# Stiffness Measurement of Retinal Capillaries and Subendothelial Matrix using Atomic Force Microscopy

**DOI:** 10.1101/2024.02.28.582372

**Authors:** Irene Santiago Tierno, Mahesh Agarwal, Nikolaos Matisioudis, Sathishkumar Chandrakumar, Kaustabh Ghosh

## Abstract

Retinal capillary degeneration is a clinical hallmark of the early stages of diabetic retinopathy (DR). Our recent studies have revealed that diabetes-induced increase in retinal capillary stiffness plays a crucial and previously unrecognized causal role in inflammation-mediated degeneration of retinal capillaries. Retinal capillary stiffening results from overexpression of lysyl oxidase, an enzyme that crosslinks and stiffens the subendothelial matrix. Since tackling DR at the early stage is expected to prevent or slow down DR progression and associated vision loss, subendothelial matrix and capillary stiffness represent relevant and novel therapeutic targets for early DR management. Further, direct measurement of retinal capillary stiffness can serve as a crucial preclinical validation step for the development of new imaging techniques for non-invasive assessment of retinal capillary stiffness in animal and human subjects. With this view in mind, we here provide a detailed protocol for the isolation and stiffness measurement of mouse retinal capillaries and retinal subendothelial matrix using atomic force microscopy.

## INTRODUCTION

Retinal capillaries are essential for maintaining retinal homeostasis and visual function. Indeed, their degeneration in early diabetes is strongly implicated in the development of vision-threatening complications of diabetic retinopathy (DR), a microvascular condition that affects nearly 40% of all individuals with diabetes ^1^. Vascular inflammation contributes significantly to retinal capillary degeneration in DR. Past studies have demonstrated an important role for aberrant molecular and biochemical cues in diabetes-induced retinal vascular inflammation ^2,3^. However, our recent work has introduced a new paradigm for DR pathogenesis that identifies retinal capillary stiffening as a crucial yet previously unrecognized determinant of retinal vascular inflammation and degeneration ^4-6^.

Specifically, the diabetes-induced increase in retinal capillary stiffness is caused by upregulation of collagen crosslinking enzyme lysyl oxidase (LOX) in retinal endothelial cells (ECs), which stiffens the subendothelial matrix (basement membrane) ^4-6^. Matrix stiffening, in turn, stiffens the overlying retinal ECs (*mechanical reciprocity*), thus leading to the overall increase in retinal capillary stiffness ^4^. Crucially, this diabetes-induced retinal capillary stiffening alone can promote retinal EC activation and inflammation-mediated EC death. This mechanical regulation of retinal EC defects can be attributed to altered endothelial mechanotransduction, the process by which mechanical cues are converted into biochemical signals to produce a biological response ^7-9^. Importantly, altered EC- and matrix-based mechanical cues have also been implicated in choroidal vascular degeneration associated with early age-related macular degeneration (AMD) ^10-12^, which attests to the broader implications of vascular mechanobiology in degenerative retinal diseases.

Notably, retinal capillary stiffening occurs early on in diabetes, which coincides with the onset of retinal inflammation. Thus, the increase in retinal capillary stiffness may serve as both a therapeutic target and an early diagnostic marker for DR. To this end, it is important to obtain reliable and direct stiffness measurements of retinal capillaries and subendothelial matrix. This can be achieved using an atomic force microscope (AFM), which offers a unique, sensitive, accurate, and reliable technique to directly measure the stiffness of cells, extracellular matrix, and tissues ^13^. An AFM applies minute (nanoNewton-level) indentation force on a sample whose stiffness determines the extent to which the indenting AFM cantilever ‘bends’-the stiffer the sample, the more the cantilever bends, and vice versa. We have used AFM extensively to measure the stiffness of cultured endothelial cells, subendothelial matrices, and isolated mouse retinal capillaries ^4-6,11,12^. These AFM stiffness measurements have helped identify endothelial mechanobiology as a key determinant of DR and AMD pathogenesis. To help broaden the scope of mechanobiology in vision research, here we provide a step-by-step guide on the use of AFM for stiffness measurements of isolated mouse retinal capillaries and subendothelial matrix.

## PROTOCOL

### 1. Isolation of mouse retinal capillaries for AFM stiffness measurement

This protocol, reported in our recent study ^4^, details the enucleation and mild fixation of mouse eye, retinal isolation and trypsin digestion, and subsequent mounting of the resultant retinal vasculature on microscopy slides for AFM stiffness measurement.

**Day 1**

#### 1.1 Enucleation and mild fixation

1.1.1. After euthanizing the mouse, insert a micro forceps behind the eyeball, hold onto the muscular attachment, and carefully pull out the eye. ***Note***: Don’t pinch the micro forces too tight or else the optic nerve may get severed.
1.1.2. Place the enucleated eye in 5% (v/v) formalin in 1X PBS for 24h at 4°C for mild fixation. ***Note***: Based on our experience ^4^, unfixed or more mildly fixed eyes yield fragile capillaries that become fragmented during AFM sample preparation and stiffness measurement. **Day 2**

#### 1.2 Retinal Isolation

1.2.1. Place the fixed eye on a piece of wax paper under a dissecting microscope.
1.2.2. Using micro forceps, hold the remnants of the muscle and optic nerve attached to the outside of the posterior eye and orient the eye so that cornea faces to one side.
1.2.3. Using a surgical blade, make an incision 1 to 2 mm behind and parallel to the limbus (cornea-sclera junction). Next, while holding the eye in place with the micro forceps, apply a downward force with the blade and continue pressing until the anterior segment of the eye is totally separated from the posterior end. Discard the anterior segment and lens. ***Note*:** Do not saw back and forth as that may cause retinal damage.
1.2.4. Transfer the posterior segment (sclera, choroid, and retina) into a 6 cm dish filled with 1X PBS.
1.2.5. Using a micro spatula, and simultaneously gently holding the optic nerve with micro forceps, scoop out the retina.
1.2.6. Store the retina in an Eppendorf tube containing 1X PBS at 4°C until capillary isolation.

#### 1.3 Retinal rinses

1.3.1. To rinse one retina, fill up six wells of a 12-well plate with 1 mL ddH_2_O. Next, using an inverted Pasteur pipette, transfer the retina from the Eppendorf tube into the **first** well
1.3.2. Rinse the retina on the orbital shaker at 120 RPM for 30 min at RT.
1.3.3. Using a P1000 pipette, carefully pipette water up and down adjacent to the retina, blowing water at the retina to cause gentle agitation.
1.3.4. Using the inverted Pasteur pipette, transfer the retina to the **second** well and repeat steps 1.3.2 and 1.3.3 four more times, with each rinse done in a fresh (ddH_2_0-containing) well.
1.3.5. Finally, transfer the retina to the <u>sixth</u> well and rinse it overnight at RT on the orbital shaker set at 100 RPM. ***Note***: The overnight rinsing facilitates separation of the retinal neuroglia from blood vessels during trypsin digestion (next step).

**Day 3**

#### 1.4 Retinal Trypsin Digest

##### Preparation Steps

1.4.1. Prepare a 10% (w/v) trypsin solution by dissolving trypsin 1:250 powder (Trypsin 1:250 Tissue Culture Grade - VWR; VWRV0458-25G) in Tris buffer (pH 8; Buffer Tris 0.1M pH 7.4 - VWR; VWRVE553-500ML). Mix gently by inverting the conical tube several times. ***Note***: Vortexing is not only unnecessary but also creates bubbles that makes it harder to confirm trypsin solubility.
1.4.2. Equilibrate the 10% trypsin solution in the 37°C bath for 10-15 min. Once the trypsin powder has completely dissolved, filter the solution using 0.2 µm syringe filter.
1.4.3. In a 12-well plate, add 2 mL/well/retina of the filtered 10% trypsin solution. Also, add some extra trypsin solution in the neighboring well (to equilibrate the Pasteur pipette walls for subsequent steps).
1.4.4. Add 3 mL ddH_2_O in three wells of a 6-well plate (dedicate these three wells of ddH_2_O for one retina).
1.4.5. In a 15 mL conical tube, insert a P200 tip before adding 4 mL of the filtered 10% trypsin solution. Transfer this conical tube in the 37°C bath to allow the tip to be soaked in trypsin for 2-3 minutes. ***Note***: Soaking the tip in trypsin helps prevent the retina from sticking to tip walls in the later steps.

##### Digestion

1.4.6. After O/N rinsing of the retina (from step 1.3.5), use a P1000 pipette to carefully pipette water up and down adjacent to the retina, squirting water on retina to cause gentle agitation.
1.4.7. Using an inverted Pasteur pipette, transfer the rinsed retina to trypsin-containing-well of the 12-well plate (from step 1.4.3). Ensure that the retina is transferred with minimal amount of residual ddH_2_O so as not to significantly dilute the trypsin solution. Incubate for 3h at 37°C
1.4.8. Next, centering the trypsin-soaked P200 tip (from step 1.4.5) on the optic nerve, gently pipette the entire vascular network up and down to dissociate the non-vascular tissue ^14^.
1.4.9. Using an inverted Pasteur pipette pre-rinsed in trypsin (by pipetting trypsin up and down five times), transfer the retinal vasculature to the <u>first</u> ddH_2_O-containing well of the 6-well plate (from step 1.4.4).
1.4.10. Swirl the plate and pipette the vasculature up and down using an inverted trypsin-rinsed Pasteur pipette to remove any residual non-vascular tissue.
1.4.11. Using the inverted Pasteur pipette, transfer the retinal vasculature to the **second** ddH2O-containing well of the 6-well plate and store at 4°C until mounting on the slide for AFM stiffness measurement. **Day 4**

#### 1.5 Mounting the Retinal Vasculature for AFM stiffness measurement

1.5.1. Using an inverted trypsin-rinsed Pasteur pipette, transfer the retinal vasculature to the **third** ddH_2_O-containing well of the 6-well plate to remove any residual trypsin solution.
1.5.2. Thoroughly clean (using ddH_2_0 and kimwipe) a Superfrost^™^ Plus microscopy slide that will be used for mounting the vascular network for AFM stiffness measurement. ***Note***: The charged surface of the slide provides strong binding for the vascular network during AFM measurement.
1.5.3. Label the slide and draw a 4-5 cm circular region around the center with a hydrophobic Aqua pen to avoid water spillage during the subsequent steps.
1.5.4. Using an inverted trypsin-rinsed Pasteur pipette, carefully transfer the retinal vasculature to the center of the marked region.
1.5.5. Using a P200 pipette, carefully remove as much excess water as possible from one edge. ***Note***: Perform this step carefully to avoid the vasculature being sucked into the tip.
1.5.6. Place the slide in a biosafety cabinet (BSL-2 hood) close to the front side (filtered mesh) and let the residual water evaporate.
1.5.7. Once the vasculature is almost dry, under a phase contrast microscope at 4x magnification, carefully rehydrate the vasculature by slowly adding ddH_2_O with a P200 pipette on to one side of the marked region. ***Note*:** Do not add ddH2O directly on the vasculature as it will detach it.
1.5.8. Take phase contrast images of the rehydrated vasculature to ensure it has good structural integrity and is properly spread out (not folding on itself) and attached to glass slide, essential criteria for reliable AFM stiffness measurement.

### 2. Obtaining subendothelial matrix from retinal microvascular endothelial cell (REC) cultures

This protocol, adapted from *Beacham DA et al*. ^13^ and reported in our recent studies ^4-6^, describes REC culture on modified glass coverslips, followed by decellularization to obtain subendothelial matrix for subsequent AFM stiffness measurement.

<u>Day 1</u>

#### 2.1 Preparation of gelatin-coated glass coverslips and cell plating

***Note***: *To avoid contamination, perform the following steps in a tissue culture hood*.

2.1.1. Place one autoclaved 12 mm dia. circular glass coverslip per well of a sterile 24-well plate.
2.1.2. Add 500 µL of pre-warmed sterile 0.2% (w/v) gelatin (diluted in PBS containing calcium and Magnesium; PBS^++^) to each well containing coverslip and incubate for 1 h at 37°C.
2.1.3. Aspirate the gelatin solution and rinse the coverslips once with sterile PBS^++^.
2.1.4. Crosslink the coated gelatin by adding 500 μL of 1% (v/v) glutaraldehyde for 30 min at room temperature (RT).
2.1.5. Collect and properly dispose (as per institutional guidelines) the glutaraldehyde solution before rinsing the coverslips with PBS^++^ for 5 min on an orbital shaker at 100 rpm in RT. Repeat this rinsing step five times. ***Note***: During the first three rinses, lift the coverslips using a sterile tweezer to allow thorough rinsing of the glutaraldehyde. Any residual amount can reduce cell viability.
2.1.6. Add 800 μL of 1M ethanolamine to the gelatin-crosslinked coverslip for 30 min at RT to quench any remaining traces of glutaraldehyde in the well.
2.1.7. Collect and properly dispose (as per institutional guidelines) the ethanolamine solution and rinse the coverslips with PBS^++^ for 5 min on an orbital shaker at 100 rpm in RT. ***Note***: During the first three rinses, lift the coverslips using a sterile tweezer to allow thorough rinsing of the ethanolamine. Any residual amount can reduce cell viability.
2.1.8. The crosslinked coverslips are now ready for cell plating. ***Note***: Although we routinely plate cells immediately after coverslip preparation, it may be worthwhile to compare cell plating on crosslinked gelatin coverslips stored in PBS^++^ at 4°C for 1-2 days.
2.1.9. Plate retinal microvascular endothelial cells (RECs) in 500 μL medium/well at a density that achieves 100% confluence within 24 h, as confirmed by phase contrast microscopy. ***Note*:** Reaching confluence within 24 h is important as the resulting REC quiescence will increase the ability of cells to secrete matrix (refer to the following step). In our experience with human RECs, a plating density of 4×10^4^ cells/cm^2^ is sufficient to reach confluence within 24 h. However, since REC size varies with species (e.g. mouse RECs are significantly smaller), the initial plating density may need to be optimized.

**Day 2 - 16**

#### 2.2 Subendothelial matrix production by HREC culture

2.2.1. After cells reach confluence (∼24 h), replace the culture medium with fresh medium supplemented with sterile-filtered ascorbic acid (final concentration of 200 ug/mL). ***Note***: Any REC treatment with disease risk factors (e.g. high glucose, advanced glycation end products etc.) or pharmacological agents can begin along with ascorbic acid treatment.
2.2.2. Change the ascorbic acid-supplemented culture medium every other day for 15 days. ***Note***: Ascorbic acid is unstable in solution. Prepare fresh 100X stock solution every other day for medium supplementation.

**End of Day 16**

#### 2.3. Decellularization of REC cultures to obtain subendothelial matrix

2.3.1. After 15 days of ascorbic acid treatment, remove the medium and rinse the cells with (calcium/magnesium-free) PBS.
2.3.2. Decellularize the REC cultures by adding 250 μL/per well of warm decellularization buffer (20 mM ammonium hydroxide and 0.5% Triton X-100 in PBS) for ∼2-3 min. Confirm the removal of cells under a phase contrast microscope.
2.3.3. Gently remove the decellularization buffer without disturbing the (REC-secreted) subendothelial matrix and add 0.5 mL of PBS to each well. ***Note***: To ensure the subendothelial matrix does not detach during this and the subsequent rinsing steps, perform gentle rinsing only with a P1000 pipette. Do not perform vacuum aspiration.
2.3.4. Store the plates at 4°C overnight to stabilize the REC-secreted matrix.

**Day 17**

#### 2.4 DNAse treatment of subendothelial matrix to remove cellular debris

2.4.1. Rinse the subendothelial matrix from step C-4 with 800 μL of PBS/well.
2.4.2. Gently remove the PBS and incubate the matrix with 200 μL of RNase-free DNase I (30 Kunitz units) at 37°C for 30 min to remove all traces of cellular debris. Check the efficiency of DNase treatment by carefully looking for cell debris under phase contrast microscope.
2.4.3. Gently remove the DNase I solution and rinse the subendothelial matrix twice with 800 μL of PBS/well.
2.4.4. The macroscale fibrous subendothelial matrix should be now visible under a phase contrast microscope. ***Note***: Visualization of the finer nanoscale matrix fibers requires AFM topographical scanning or high-resolution confocal imaging of immunolabeled matrix proteins.
2.4.5. Immediately use the fresh (unfixed) subendothelial matrix for AFM stiffness measurement.
2.4.6. Following AFM measurement, matrix samples can be fixed with 1% (v/v) paraformaldehyde (15 min at room temperature) and stored at 4°C for immunolabeling with antibodies against subendothelial matrix proteins (e.g. collagen IV) or matrix modulating factors (e.g. lysyl oxidase).

### 3. AFM stiffness measurement

This protocol, adapted from the Bruker NanoWizard® Series user manual and reported in our recent studies ^4,5^, details the acquisition and analysis of AFM stiffness data from retinal capillaries and subendothelial matrix using Bruker’s NanoWizard® 4 XP Bioscience AFM. Although the steps outlined below are based on this specific model of AFM, the underlying principles are generally applicable to all AFM models. AFM measurements are performed with a small probe attached to the tip of a sensitive cantilever that is lowered to indent the sample up to a preset indentation force, before being raised to its original height. Sample stiffness is calculated from the cantilever deflection during indentation.

#### 3.1 Cantilever probe selection

***Note***: Selection of appropriate cantilever stiffness (spring constant, k) and probe dimension is essential for reliable and sensitive measurements. Specifically, soft biological samples require soft cantilevers that can bend upon sample indentation while the probe dimensions are selected to match the dimensions of the sample.

3.1.1. For stiffness measurement of 5-8 µm diameter mouse retinal vessels, we use the Bruker SAA-SPH 1 µm radius cantilever probe with a spring constant (k) of 0.2-0.3 N/m.
3.1.2. For stiffness measurement of subendothelial matrix composed of nanoscale fibers, we use the Bruker PFQNM-LC-A 70 nm radius cantilever probe with a spring constant (k) of 0.06-0.1 N/m.

#### 3.2. Cantilever mounting

3.2.1. On the AFM computer, open the software that control the AFM unit (JPK SPM) and the camera attached to a phase contrast microscope that is used to visualize the cantilever and sample.
3.2.2. Fix the cantilever holder onto the cantilever changing stand and mount the selected cantilever on the holder using watchmaker forceps under stereoscope without physically damaging the cantilever. Tighten the screw on the holder to secure the cantilever.
3.2.3. Place and lock the cantilever holder on the AFM head that is resting on the stand.
3.2.4. Using the step motor function in the AFM software, withdraw the AFM head to the highest point and set that position as ‘point zero’ in the software. ***Note***: This step prevents the AFM cantilever from accidentally hitting the sample stage while mounting the AFM head (next step).
3.2.5. Carefully lift the AFM head from its stand and mount it on the AFM sample stage by placing the legs in their respective slots.

#### 3.3. Laser alignment

***Note***: Focusing the infrared laser beam on the tip of cantilever and centering the reflected laser spot on the photodetector are essential for precise detection of cantilever deflection and, consequently, stiffness measurement. The initial laser alignment is performed without any sample (i.e., in ‘Air’ mode) as it ensures a more precise alignment of laser beam on the cantilever tip (by preventing laser refraction in liquid). However, as biological samples are immersed in a medium or buffer that has a different refractive index than air, it is important to realign the laser before measuring sample stiffness.

3.3.1. For **laser alignment in air**, place the AFM head on the sample stage and use the 10X objective and attached camera to visualize the cantilever on the monitor in live mode. ***Note***: The infrared laser cannot be seen by naked eye but is visible with CCD camera.
3.3.2. Select ‘contact mode force spectroscopy’ function on the software and open the window entitled ‘Laser alignment’.
3.3.3. Using the two screws marked ‘laser lateral’ on the AFM head, focus the laser beam on the cantilever such that the ‘SUM’ value in the laser alignment window is the highest. ***Note***: Typically, the highest value for SUM is obtained by focusing the laser at or around the center of the cantilever.
3.3.4. Using the “Detector adjustment screws”, adjust the photodetector such that the laser spot is at the center of the laser alignment window.
3.3.5. Using the ‘Mirror adjustment screw’, adjust the mirror to ensure that the ‘SUM’ value is at the highest possible value. ***Note*:** A high SUM value ensures sensitive and accurate assessment of sample stiffness. Thus, both the laser alignment and mirror adjustment steps should be performed to obtain the highest SUM value.
3.3.6. Next, for **laser alignment in liquid**, add the desired culture medium or buffer in a dish and place it on the stage. ***Note*:** The dish should be mounted firmly on stage to prevent any vibration or drift, which create measurement artefacts during sample indentation.
3.3.7. While looking at the live camera feed focused on the cantilever, lower it into the liquid using the step motor function.
3.3.8. Repeat steps C-4 and C-5 to ensure that the SUM value remains the highest and the laser spot is at the center of the laser alignment window.

#### 3.4. Calibration of cantilever spring constant

Although cantilever probes come with a manufacturer-calibrated spring constant(k), it is good practice to independently verify its value (in liquid) before the start of a measurement. Cantilevers must be calibrated in liquid and on a clean glass surface that minimizes unwanted attractive or repulsive interactions between the cantilever probe and glass surface.

3.4.1. Set the setpoint force applied by the cantilever at 1.5 V ***Note***: Setpoint force sets the maximum laser deflection from cantilever that is allowed for a force indentation. The setpoint should be greater than the resting vertical deflection of a cantilever that results from its thermal vibration or electrostatic interaction. If the cantilever fails to ‘Approach’ at a given setpoint force, increase it incrementally until it indents the sample and generates a force indentation curve.
3.4.2. For calibration of Spring Constant on glass, set Z Length at 1 µm. ***Note*:** Z Length signifies the maximum distance by which the z-piezo withdraws after cantilever has reached the setpoint during sample indentation. Z length should be at least enough to ensure the cantilever probe separates cleanly from the sample but not too much to slow down multiple force indentation measurements at a location during contact force spectroscopy mode or high-speed QI force mapping.
3.4.3. Set Z speed at 2 µm/s. ***Note*:** Z speed refers to the speed at which the z-piezo moves the cantilever vertically down towards the sample during force indentation. It should be optimized because a very low speed with primarily capture the viscous behavior of the sample while a very high speed will primarily capture the elastic behavior. The optimal speed is expected to capture the true viscoelastic nature of biological samples.
3.4.4. Set target height on the Z range at 7.5 µm. ***Note*:** Target height on the Z range indicates the approximate distance from the sample at which the z-piezo rests after measuring a force curve and before moving to a different location on the sample. The Z Range target height should be greater than the typical height of the sample features.
3.4.5. Click ‘Approach’ in the ‘contact mode force spectroscopy’ window. ***Note*:** Once the step motors have brought the cantilever fairly close to the sample surface (judging by the focus in the phase contrast image), use the ‘approach’ feature to bring down the cantilever in smaller increments of 15 µm.
3.4.6. Click ‘Acquire’ to capture a force curve. ***Note*:** Cantilever deflects during sample indentation, with the z-piezo (driving the cantilever vertically) retracting once the vertical (laser) deflection reaches the setpoint force. During probe retraction from the sample, any adhesion between these surfaces may cause a reverse cantilever deflection, causing the retraction curve to dive below the baseline. The cantilever will eventually rest at the Z target height set at step D-4.
3.4.7. Select the force curve, open it in the calibration manager window, and select ‘contact-based mode’ in the method section.
3.4.8. Using ‘Select fit range’ function, select the cantilever ‘retraction’ curve for a *linear curve* fit. Next, check the ‘sensitivity check box’ to convert the force unit from Volt to Newton.
3.4.9. Lift the cantilever by 100-200 µm in the liquid and select the ‘Thermal noise’ function. Again using ‘Select fit range’, fit the thermal noise bell curve with a Lorentz curve. After curve fitting, select the spring constant(k) box. Confirm that spring constant (k) is close to the manufacturer’s value and note it down for future reference. ***Note*:** Thermal noise is the natural frequency of the cantilever at a particular temperature. Correction for thermal noise is required for accurate spring constant measurement.
3.4.10. After calibration, the unit for setpoint will change from m-Volt to nano-Newton.

#### 3.5. Acquisition of force-distance curves

3.5.1. Place the slide-mounted sample (retinal vessels or subendothelial matrix) on the AFM stage.
3.5.2. Select ‘contact mode force spectroscopy’ in the JPK software experiment section.
3.5.3. Set the setpoint force at 0.5 nN and click ‘Approach’. ***Note***: This setpoint is significantly higher than the thermal deflection of both SAA-SPH 1 µm & PFQNM-LC-A 70 nm cantilever probes, which ensures a smooth cantilever approach towards the sample.
3.5.4. After the cantilever has detected the surface and returned to its resting ‘target height’, set the setpoint at 0.2 nN for the stiffer SAA-SPH 1 µm cantilever probe or 0.1 nN for the softer PFQNM-LC-A 70 nm cantilever probe. ***Note***: The set point should be adjusted based on both cantilever and sample type. Stiffer samples and cantilevers require higher setpoint because they do not indent/bend readily, and vice versa for softer samples and cantilevers (refer to steps A-1,2).
3.5.5. Set Z Length around 2.5 µm as it ensures clean separation of the cantilever probe from viscous biological samples during retraction (refer to step D-2).
3.5.6. Set Z speed at 2 µm/sec (refer to step D-3).
3.5.7. Using the stage screws on the AFM stage, carefully position the cantilever probe at a desired location on the sample. ***Note*:** Since capillaries do not always spread flat on the slide, it is safe to withdraw the cantilever a further 10-20 µm from its resting height before moving the stage.
3.5.8. Click ‘Approach’ to first bring the cantilever closer to sample and then click ‘Acquire’ to capture the force curves for the desired locations. Save all force curves for analysis. ***Note***: Biological samples like cells, matrix, and blood vessels are viscoelastic by nature and thus may undergo some permanent deformation and/or change in apparent stiffness following force indentation. If so, this will be reflected in the misalignment of approach and retraction curves (hysteresis).

#### 3.6. Data analysis

3.6.1. Open the force curve in the JPK data processing software.
3.6.2. Select the ‘retraction’ force curve for data analysis (similar to step D-8).
3.6.3. Using ‘Data smoothing’ function, smoothen the force curve with gaussian filter to remove the unwanted noise in the acquired data.
3.6.4. Using ‘Baseline subtraction’ function, adjust the value of the slope and magnitude of the (non-contact) baseline portion of the force curve to zero.
3.6.5. Next, click on the ‘contact point’ function to automatically bring the contact point of the force curve to the (0,0) coordinates on X- and Y-axes.
3.6.6. Then, using ‘Vertical tip position’ function, calculate the actual vertical position of the cantilever on the Y-axis by correcting for any sample indentation.
3.6.7. On the processed force curve, apply the ‘elasticity fit’ function by first selecting the tip shape and radius (based on the selected cantilever probe), followed by force curve fitting using the Hertz/Sneddon model. ***Note***: If matrix indentation by the 70 nm radius probe exceeds 70 nm, which is typically the case, select the ‘paraboloid’ tip shape. For capillary stiffness measurement, sample indentation by the 1 um radius probe never exceeds 1 um, so select the ‘sphere’ tip shape. Further, if the contact point of fitted curve does not coincide with the contact point of actual retraction curve, select the ‘shift curve’ check box.
3.6.8. Note and save the value of Young’s modulus (stiffness).

## REPRESENTATIVE RESULTS

### Mouse retinal capillaries

AFM stiffness measurement of isolated retinal capillaries involves sample handling steps that could potentially damage their mechanostructural integrity. To prevent this and thereby ensure the feasibility, reliability, and reproducibility of AFM measurements, the enucleated eyes are fixed in 5% formalin overnight at 4°C prior to vessel isolation. This mild fixation protocol with reduced formalin concentration, low fixation temperature, limited fixation time, and lack of corneal puncture was developed to minimize any potential crosslinking/stiffening artefacts caused by chemical fixation. As shown in **Fig. 1**, this relatively mild fixation ensures that the isolated retinal vasculature is structurally robust and sufficiently durable for the AFM measurement. In contrast, retinal vessels isolated from unfixed eyes (using hypotonic method) or briefly fixed eyes (for 8h) become fragmented or collapse, thereby making them unsuitable for AFM measurement **(Fig. 1)**.

**Fig. 1:**
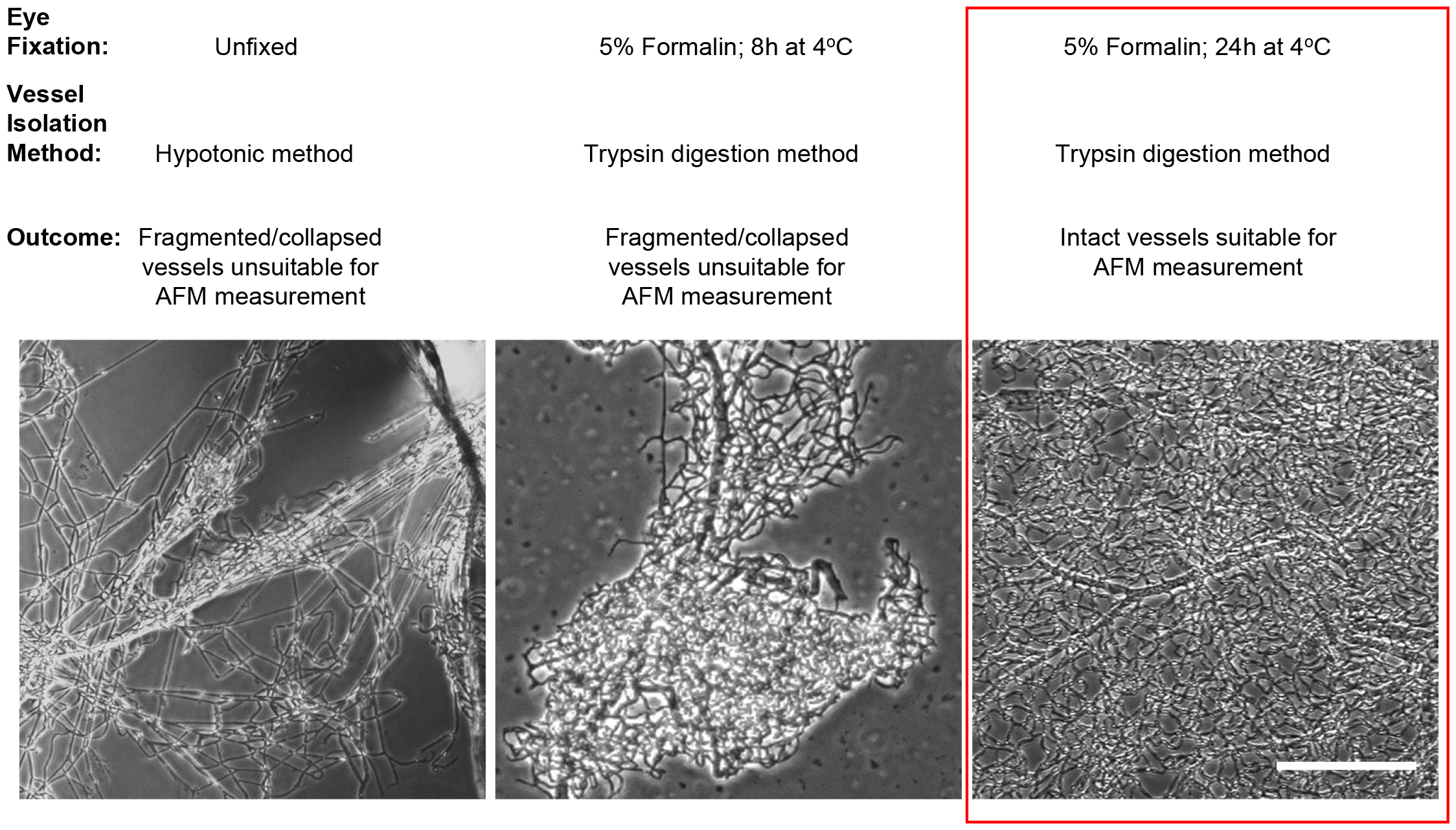
Protocol optimization for isolation of intact mouse retinal vessels for AFM measurement. Representative phase contrast images show mouse retinal vessels obtained using the different isolation methods. Comparing the structural integrity and durability of the isolated vessels, trypsin digestion of retinas from enucleated eyes fixed in 5% formalin for 24h at 4°C (red box) was found to yield the most suitable retinal vasculature for AFM stiffness measurement. Retinal vessels isolated using this method exhibited nice vascular network that spread uniformly along the glass surface. Scale bar: 500 um. Figure reproduced from Chandrakumar S et al. *Diabetes* (2024); 73: 280-291 © American Diabetes Association

### Retinal subendothelial matrix

Vascular stiffness reflects the combined stiffness of vascular cells and the basement membrane (subendothelial matrix) ^4^. Since cells adapt to matrix stiffness by undergoing a similar change in their own stiffness, a process termed ‘mechanical reciprocity’ ^9^, subendothelial matrix stiffness becomes an important determinant of the overall vascular stiffness. For matrix stiffness measurement, it is important to obtain a homogeneously dense subendothelial matrix. For human retinal ECs grown in ascorbic acid-supplemented culture medium, this usually takes 10-15 days **(Fig. 2A)** ^4-6^. This difference in culture period may arise from lot-to-lot differences in commercially available primary retinal ECs. Further, we generally find that commercially available retinal ECs from C57BL/6 mice deposit a denser matrix when compared with primary human retinal EC culture, thus indicating species-specific differences. As shown in **Fig. 2A**, phase contrast images only provide a gross view of the matrix at a macro scale. However, the finer nano-to-micro-scale fibrillar structure becomes apparent in high resolution confocal fluorescence images of the matrix immunolabeled with antibodies against matrix structural proteins collagen IV and fibronectin **(Fig. 2B)**. It should be noted that these matrix proteins also provide instructive cues to endothelial cells by binding to specific integrin receptors ^9^.

**Fig. 2:**
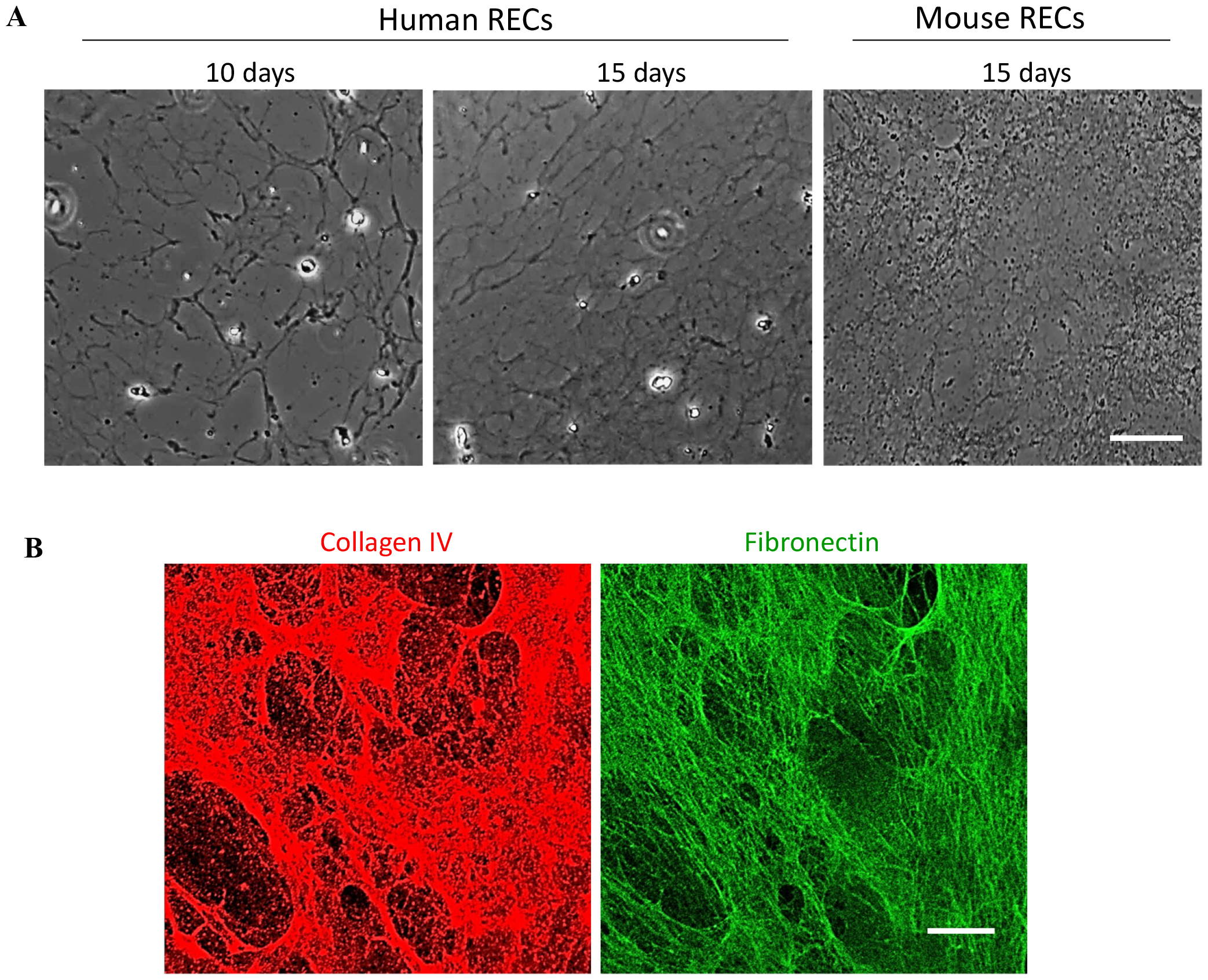
Decellularized matrices obtained from primary human and mouse REC cultures. **(A)**Representative phase contrast images show subendothelial matrix aggregates on glass coverslips following decellularization of 10d or 15d cultures of human or mouse RECs. Scale bar: 100 um. **(B)**Representative high magnification confocal fluorescence images of decellularized matrices obtained from 15 d human REC cultures and labeled with anti-collagen IV and anti-fibronectin antibodies reveal a dense nanofibrillar collagen IV and fibronectin matrix. Scale bar: 20 um.

### AFM stiffness measurement

An AFM stiffness measurement begins with the z-piezo moving the cantilever vertically down towards the sample. There is no cantilever deflection at this time, which produces a flat baseline of the approach curve **(Fig. 3A)**. As the cantilever probe contacts and indents the sample, the cantilever bends, causing laser deflection on the photodetector, which is depicted by the vertical deflection of the approach curve. After applying a preset indentation force on the sample, the cantilever retracts to the starting position away from the sample. The deflected retraction curve is then fitted to the Hertz/Sneddon model to calculate the sample’s Young’s modulus (stiffness). From the representative force indentation measurement shown in **Fig. 3**, it is clear that the approach and retraction curves obtained from a retinal capillary isolated from diabetic mouse **(Fig. 3B)** are substantially steeper than those obtained from a nondiabetic mouse **(Fig. 3A)**. Steeper slope of force indentation curves indicates greater cantilever deflection caused by higher sample resistance to force indentation, which reflects higher sample stiffness ^13^. Indeed, subsequent data analysis revealed that mouse retinal capillaries become significantly stiffer in diabetes ^4^. It should also be noted that contact between the cantilever probe and biological samples often causes nonspecific surface adhesion, which leads to negative cantilever deflection during retraction, as seen from the extension of retraction curve beyond the baseline **(Fig. 3B)**. As we have previously reported, stiffness (Young’s modulus) of subendothelial matrices obtained from retinal EC cultures is also calculated in the aforementioned manner ^5,6^.

**Fig. 3:**
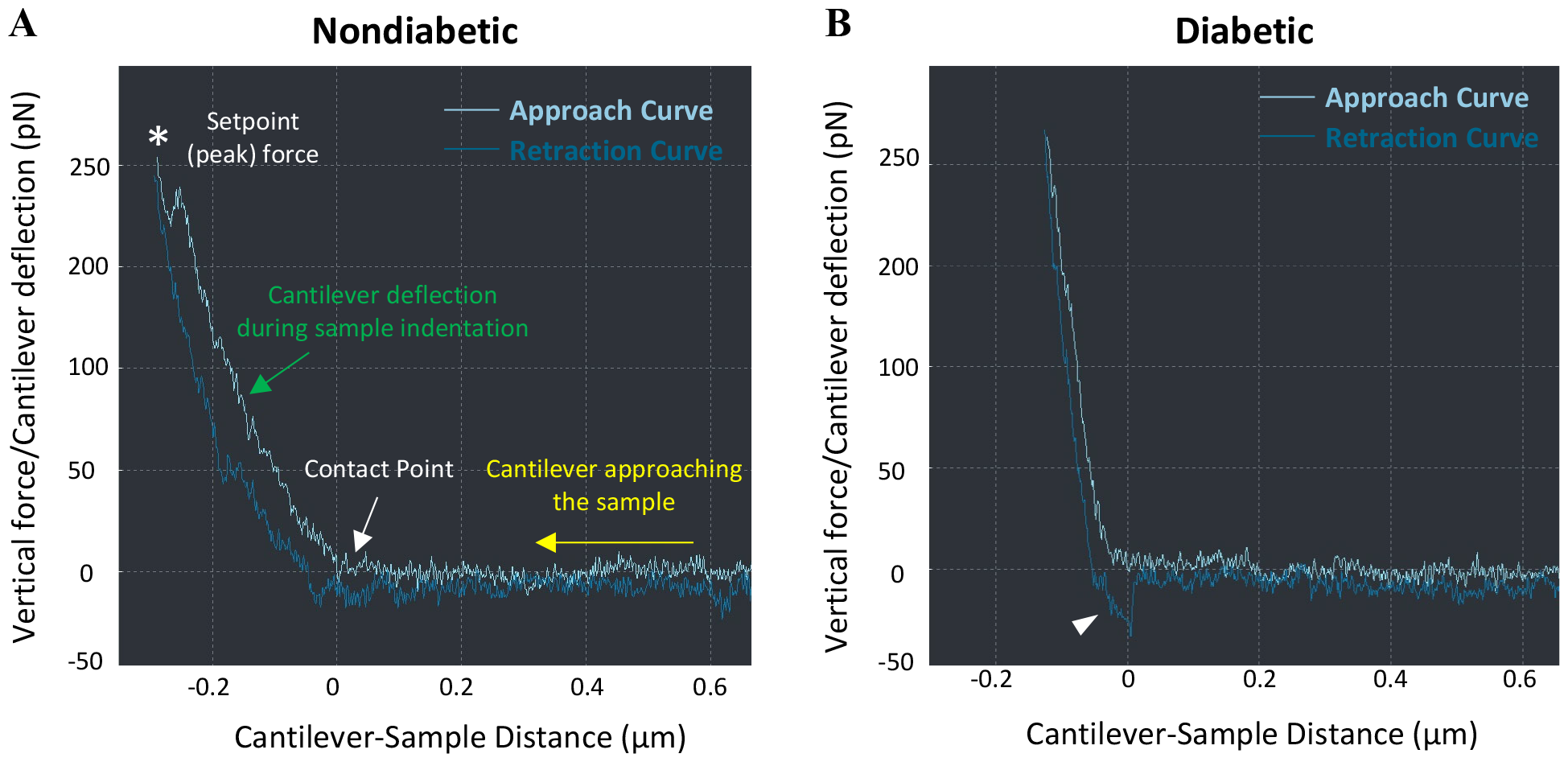
Force distance curves from AFM stiffness measurement of mouse retinal capillaries. Line graphs indicate a representative ‘approach’ (light color) and ‘retraction’ (dark color) curve from a single force indentation measurement at one location of a mouse retinal capillary isolated from a nondiabetic **(A)** or diabetic **(B)** mouse. The force curves, obtained using a SAA-SPH 1 µm radius hemispherical cantilever probe, plot the relationship between cantilever-sample distance (controlled by the z-piezo) and the applied vertical force that causes cantilever deflection. **(A)** Yellow arrow indicates the z-piezo-driven cantilever approach towards the sample, white arrow indicates the “contact point” where the cantilever probe makes contact with the sample, and the green arrow indicates the cantilever deflection up to a setpoint (peak) indentation force (*). **(B)** Both approach and retraction curves obtained from a retinal capillary isolated from diabetic mouse exhibit a markedly steeper slope than those from their nondiabetic counterparts (shown in A), which indicates higher capillary stiffness in diabetic mice. Arrowhead indicates a dip in the retraction curve below baseline, which reflects negative deflection of the cantilever probe caused by adhesion between the probe and sample during indentation.

## DISCUSSION

AFM has been widely used to measure disease-associated changes in the stiffness of larger vessels such as aorta and arteries ^15^. These findings have helped establish a role of endothelial mechanobiology in cardiovascular complications such as atherosclerosis ^16^. Based on these findings, we have begun to investigate the previously unrecognized role of endothelial mechanobiology in the development of retinal microvascular lesions in early DR. Success in this pursuit, however, relies on the accurate measurement of diabetes-induced changes in retinal capillary stiffness. Our recent studies have revealed that similar to large vessels, stiffness of retinal capillaries and retinal EC-secreted subendothelial matrix can also be directly, accurately, and reliably measured using AFM ^4-6^.

Our approach involves the isolation of mouse retinal capillaries from mildly fixed eyes using trypsin digestion. Based on our experience, this mild fixation step is necessary as capillaries isolated from unfixed eyes (using hypotonic method) are fragile and become fragmented, rendering them unsuitable for AFM stiffness measurement ^4^. As an alternative approach, a recent study reported AFM stiffness measurement of retinal capillaries from lightly fixed retinal flat mounts ^17^. By performing AFM force indentations on retinal capillaries within the intact retina, this approach enables ‘in situ’ stiffness measurement. However, these capillary stiffness measurements likely include the stiffness of inner limiting membrane. Further, this approach is restricted to the measurement of only superficial capillaries that are accessible on a retinal flat mount, while leaving out the deeper-lying capillary plexus that is specifically affected in early DR ^18^. These issues can be addressed using our approach that extracts vessels cleanly (devoid of residual retinal tissue) from all retinal layers. That said, we realize that the mild formalin fixation employed in our protocol may cause a crosslinking-associated stiffening artefact in our AFM measurements. Therefore, it would be prudent to interpret the stiffness values obtained using our approach more in terms of relative changes in stiffness (between different experimental groups) rather than absolute stiffness.

In contrast to retinal capillaries that require mild fixation, subendothelial matrix obtained from retinal EC cultures can be used without any modification. This is largely due to the robustness of deposited matrix, which results from a combination of several factors including the addition of ascorbic acid in culture medium (which enhances collagen synthesis), chemical modification of glass coverslips that prevents matrix detachment during decellularization, and prolonged duration of culture (10-15 days). Importantly, we have shown that the hyperglycemia-induced increase in subendothelial matrix stiffness *in vitro* is consistent with the diabetes-induced increase in retinal capillary stiffness seen *in vivo* ^4^. This is not surprising as an increase in matrix stiffness is expected to increase the stiffness of the overlying retinal ECs (due to mechanical reciprocity) that, together, increase the overall capillary stiffness. Indeed, we recently showed that retinal ECs become stiffer when grown under diabetic conditions ^4^, likely via an increase in Rho/ROCK-dependent actin cytoskeletal tension ^19^. Still, the fact that stiffness alterations in unfixed subendothelial matrix mirrors the trend seen in mildly fixed intact retinal vessels testifies to the validity of our mild fixation approach.

Direct stiffness measurement of retinal capillaries from animal models of DR not only helps establish endothelial mechanobiology as a novel therapeutic target for DR management, it can also serve as a crucial preclinical validation step for the development of imaging techniques for non-invasive assessment of retinal capillary stiffness in DR patients. AFM stiffness measurements of retinal capillaries isolated from post-mortem human donor eyes will provide important proof-of-concept in this regard. Given that retinal capillary stiffness increases early on in the (streptozotocin) mouse model of type 1 diabetes ^4^, successful use of AFM in validating stiffness-measuring imaging modalities may lead to the identification of retinal capillary stiffness as a clinical biomarker for early DR pathogenesis.

## ACKNOWLEDGEMENTS

This work was supported by National Eye Institute/NIH grant R01EY028242 (to K.G.), The Stephen Ryan Initiative for Macular Research (RIMR) Special Grant from W.M. Keck Foundation (to Doheny Eye Institute), and Ursula Mandel Fellowship and UCLA Graduate Council Diversity Fellowship (to I.S.T.). This work was also supported by an Unrestricted Grant from Research to Prevent Blindness, Inc. to the Department of Ophthalmology at UCLA.

